# LIPC-E97G Gain-of-function variant potentiates the hypocholesterolemic and anti-atherogenic effects of evinacumab in *Ldlr*^−/−^ mice

**DOI:** 10.64898/2026.09.28.754893

**Authors:** Lucie Vince, Amélie Thouzeau, Audrey Ayer, Bertrand Cariou, Antoine Rimbert, Cédric Le May

## Abstract

**Introduction:** Cardiovascular disease is the leading cause of death worldwide (Dagenais et al. 2020), and its main risk factor is LDL-C (Hagenbeek et al. 2020). Only a few genes are associated with LDL-C concentrations, and only 2 genes, *ANGPTL3* (Musunuru et al. 2010) and *LIPC*, are associated with familial combined hypocholesterolemia. Our team recently identified the second genetic cause associated with familial combined hypocholesterolemia, a gain-of-function mutant of the *LIPC* gene encoding hepatic lipase (HL), *LIPC*-E97G (Dijk et al. 2022). Although the mode of action of this variant remains largely unknown, we have demonstrated that it significantly increases the phospholipase activity of HL and affects lipoprotein metabolism by promoting plasma cholesterol clearance, particularly of LDL and HDL.

**Objective:** The aim of this study was to explore the additive effect of our HL-E97G variant with a pharmacological inhibitor of the ANGPTL3 protein, evinacumab, on cholesterolemia and atherosclerosis.

**Materiel and methods:** Atherosclerosis measurements were performed on *Ldlr*^−/−^ mice fed a pro-atherogenic diet. These mice received a single injection of a liver-targeting AAV8 expressing the wild-type or E97G human form of the *LIPC* gene (1×10^11^ genome copies/mouse) and a weekly injection of NaCl or evinacumab (25 mg/kg/week).

**Results:** From a therapeutic approach, we demonstrated a cumulative effect of HL-E97G with evinacumab. Atherosclerotic lesions in the aortic arch of *Ldlr*^−/−^ mice were reduced by 36% (p=0.4897) with evinacumab, by 65% (p-value=0.0028) with HL-E97G, and by 80% (p-value < 0.0001) with the combined therapy.

**Conclusion:** Our data demonstrate that HL-E97G reduces the development of atherosclerosis in mice in an LDLR- and ANGPTL3-independent manner, strongly reinforcing the therapeutic potential of HL-E97G for patients at high risk of cardiovascular disease.

## Introduction

Cardiovascular disease is the leading cause of death worldwide, representing 40% of all deaths (Dagenais et al. 2020). Atherosclerotic cardiovascular disease (ASCVD) is a major component of cardiovascular disease. 76.9% of patients at high and very high cardiovascular risk undergoing lipid-lowering therapy have atherosclerosis (Ray et al. 2023). It is characterized by a deposit of lipids in the arteries, also known as atheroma, leading to their long-term obstruction and ischemia. Consequently, one of its main risk factors is high concentrations of LDL-cholesterol (LDL-C), and it has a high heritability of plasma cholesterol levels (Hagenbeek et al. 2020; Borén et al. 2020). Despite existing treatments (statins, ezetimibe, PCSK9 inhibitors, lomitapide, evinacumab…), 80% of patients fail to reach their LDL-C goals (Ray et al. 2023), leaving them with poor protection against cardiovascular disease, particularly for patients with familial hypercholesterolemia (FH).

FH is a genetic disease affecting one person in 311 (Beheshti et al. 2020). It is characterized by a high concentration of LDL-C due to a genetic mutation in the *LDLR, APOB*, and *PCSK9* genes, leading to hypercholesterolemia. In the most severe cases, homozygous familial hypercholesterolemia (HoFH), the LDL receptor (LDLR) no longer functions due to a biallelic mutation of the *LDLR*. This condition is rarer, affecting one person in 400,000 (Beheshti et al. 2020). Because lipid-lowering treatments are LDLR-dependent, it is difficult to treat HoFH patients.

More recently, a new drug has been brought to market: evinacumab, an ANGPTL3 inhibitor. *ANGPTL3* loss-of-function variants have been shown, for the first time, to be associated with familial combined hypocholesterolemia, i.e., low levels of LDL-C and HDL-C (Musunuru et al. 2010). This new treatment has an LDLR-independent mode of action, reducing LDL-C levels by 56% in HoFH patients (Béliard et al. 2024).

Our team has recently identified the second genetic cause associated with familial combined hypocholesterolemia: the *LIPC*-E97G variant (Dijk et al. 2022). The *LIPC* gene encodes hepatic lipase (HL), a protein on the surface of hepatocytes involved in the hydrolysis and internalization of lipoproteins. This mutated form of HL displays triglyceride lipase activity similar to that of the wild-type, but the phospholipase activity is higher. HL-E97G is characterized by a very high cholesterol clearance of lipoproteins and a strong anti-atherogenic effect, LDLR-independent (Sotin et al. 2025). These results reinforce the therapeutic potential of HL-E97G, particularly for HoFH patients. We therefore wanted to know whether the hypocholesterolemic and anti-atherogenic effects of HL-E97G could reinforce the effects of evinacumab, an ANGPTL3 inhibitor.

## Methods

### Mouse study

*Ldlr*-deficient (*Ldlr*^−/−^) mice (C57Bl/6J background) were bred at the University of Nantes animal facility (Nantes, France) and used for experiments approved by the Pays de la Loire ethics committee (France, 006) and the Ministry of Higher Education and Research (France). All mice were housed in groups under a 12-hour day-night cycle at 22°C and had free access to food and water.

Male (10-27 weeks old) and female (8-18 weeks old) *Ldlr*^−/−^ mice were randomized into 4 blocks (n = 4-6 for males and n = 8-9 for females) based on age, body weight, and plasma cholesterol, using the “blockrand” function on RStudio, and are fed a pro-atherogenic diet containing regular casein, 1.25% cholesterol, and 0.5% sodium cholate (Research Diet #D12109Ci). The mice then receive a single intravenous injection containing 1×10^11^ genome copies of the AAV8 expressing, under the TBG (liver-specific) promoter, the WT or E97G form of hepatic lipase (HL-WT or HL-E97G) (Vector Biolabs, USA). And every week, the mice were injected subcutaneously with evinacumab (EVKEEZA), a human anti-ANGPTL3 antibody, at 25 mg/kg/week.

### Body weight and plasma lipids

Mice are weighed weekly. Blood is collected every two weeks from the tail of the mouse after a 3-hour fast (gastric emptying from 11 am to 2 pm) using a heparinized capillary (Hirschmann #9100260), and plasma is harvested after centrifugation (4°C, 5 minutes, 6000 RPM). Plasma cholesterol (DiaSys #113009910026) and triglycerides (Human #10725) are measured by colorimetric assay.

### Quantification of atherosclerosis

12 weeks after virus injection, mice are anesthetized with an intraperitoneal injection of ketamine/xylazine (100 mg and 20 mg/kg, respectively) followed by cardiac exsanguination and PBS perfusion. The aortic arch and root were dissected under a binocular loupe, embedded in OCT (Sakura #4583), and stored at −80°C. Cross-sections of the aortic arch and root, 5µm thick, were then taken at −20°C on a CryoStar NX70 cryostat (Thermo Fisher Scientific, USA), embedded in SuperFrost® Plus slides (Thermo Fisher Scientific, USA), and stored at −80°C (Bessueille et al. 2023). Tissues are then stained with Oil Red O (Sigma-Aldrich #O0625) (Mehlem et al. 2013) and scanned with the NanoZoomer S360 (Hamamatsu #C13220-01) within 24 hours. Atherosclerotic plaque area is quantified with NDP.view2 software (Hamamatsu #U12388-01).

### Statistics

Data were analyzed using GraphPad Prism 10 software. A two-way ANOVA test, with multiple comparisons, was used for the analysis of body weight and plasma cholesterol. A one-way ANOVA test, non-parametric and multiple comparison, was applied for the analysis of atherosclerosis in the aortic root and aortic arch. Data are visualized with mean ± SEM, and P > 0.05 is significant.

## Results

### HL-E97G drastically reduces plasma cholesterol levels compared with ANGPTL3 inhibition in *Ldlr*^−/−^ mice

To test whether the hypocholesterolemic effect of ANGPTL3 inhibition by HL-E97G is reinforced, we measured plasma cholesterol over 12 weeks in dyslipidemic *Ldlr*^−/−^ mice expressing HL-WT or -E97G and weekly injected with NaCl or the anti-ANGPTL3 antibody evinacumab (Figure 1A). Expression of HL-E97G and evinacumab had no impact on body weight (Figure 1B). Evinacumab decreased plasma cholesterol by 29% (p-value = 0.2024, Figure 1C). However, HL-E97G is drastically more effective, reducing plasma cholesterol by 79% (p-value < 0.0001, Figure 1C). The hypocholesterolemic effect of HL-E97G is so strong that the effect of HL-E97G/evinacumab co-treatment is not very marked (−5%, p-value = 0.7087, Figure 1C) but still reduces cholesterol levels by 84% (p-value < 0.0001, Figure 1C).

**Figure 1.**
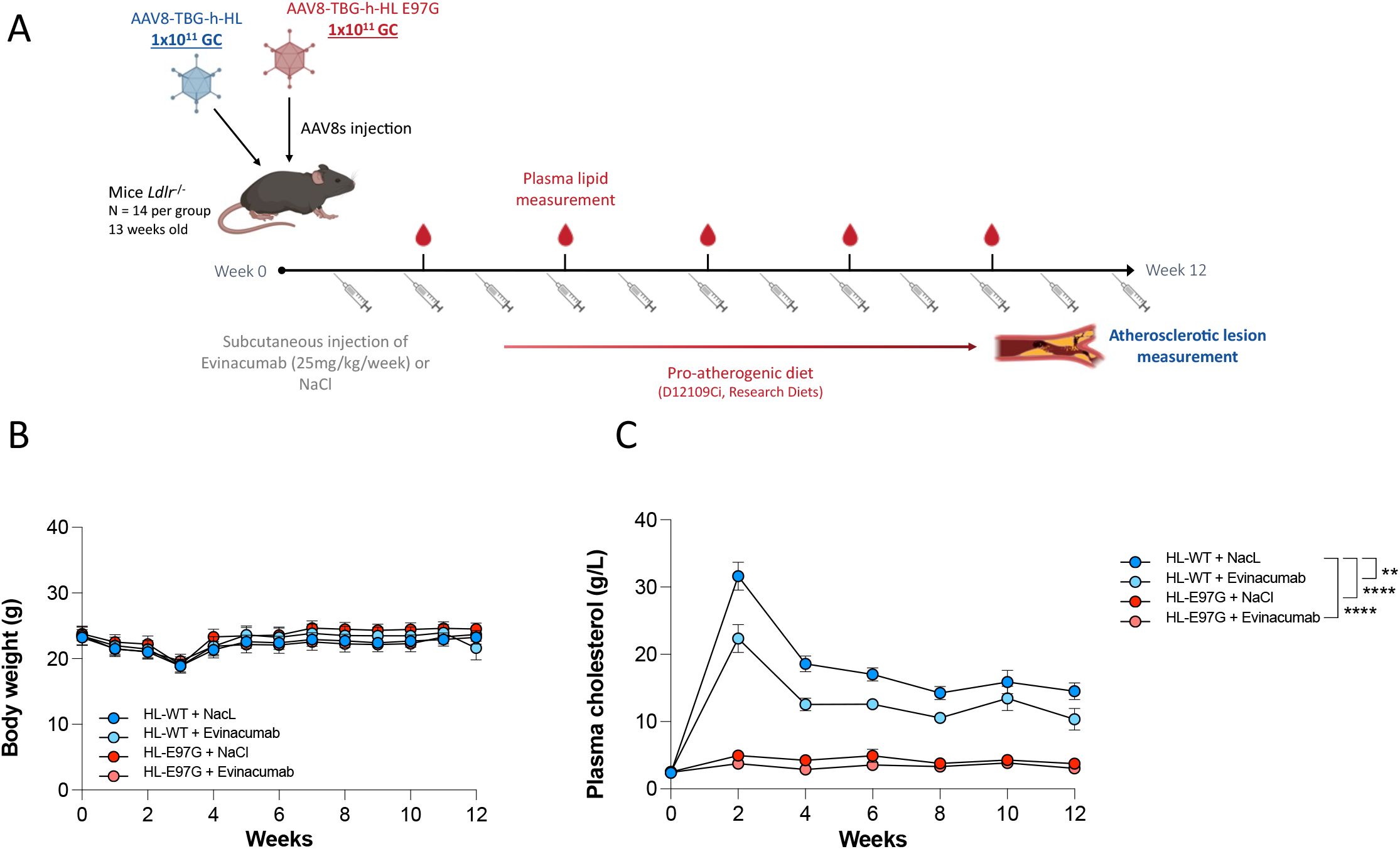
HL-E97G expression drastically decreases plasma cholesterol levels compared to Evinacumab alone. (A) Schematic diagram of experimental protocol. (B) Body weight and (C) plasma cholesterol are measured according to the above protocol. A Two-way ANOVA test, with multiple comparison, was used. Data are visualized with mean ± SEM and P > 0.05 is significant. * p < 0.05, ** p< 0.01, *** p < 0.001, **** < 0.0001.

### HL-E97G/evinacumab co-treatment has a strong additive effect and a potent anti-atherogenic effect

To determine the anti-atherogenic properties of HL-E97G and ANGPTL3, aortic roots and arches were harvested 12 weeks after virus injection (Figure 1A). Evinacumab tended to limit the development of atherosclerosis by 14% in the aortic root (p-value = 0.3487, Figure 2A). In contrast, HL-E97G significantly reduced the progression of atherosclerosis by 24% compared to HL-WT + NaCl control mice (p-value = 0.0398, Figure 2A). The anti-atherogenic effect of HL-E97G/evinacumab co-treatment was very strong −32% compared to control mice (p-value = 0.0004, Figure 2A).

**Figure 2.**
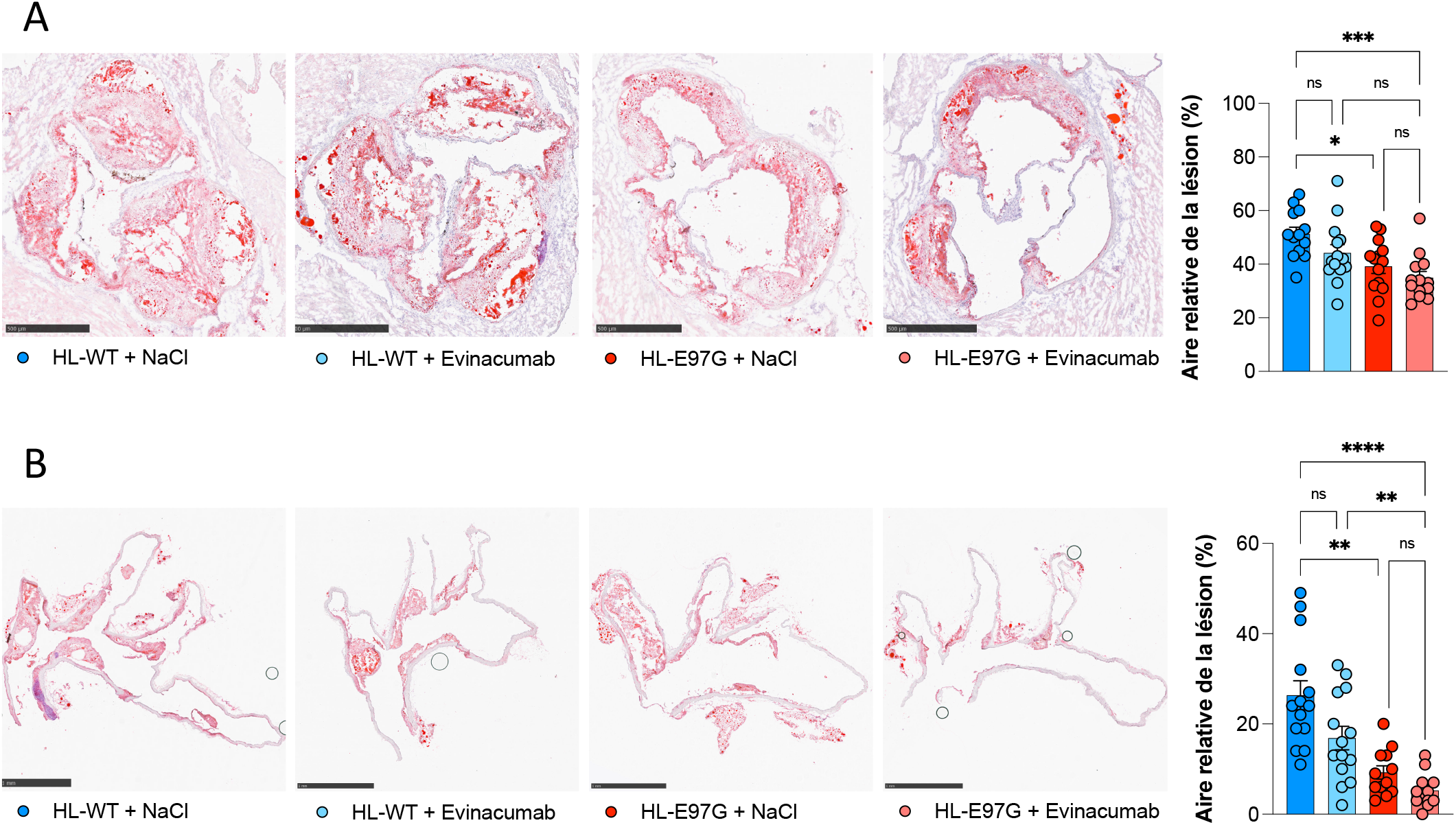
HL-E97G and Evinacumab co-treatment showed a strong anti-atherogenic effect. (A) Transverse sections of the root and (B) sagittal sections of the aortic arch, stained with oil red o, enabling atheromatous plaques to be quantified. A non-parametric, multiple comparison, One-way ANOVA test was applied. Data are visualized with mean ± SEM and P > 0.05 is significant.

Atherosclerosis was also quantified in the aortic arch, where atherosclerotic plaques develop more slowly. The ANGPTL3 inhibitor tended to limit the development of atherosclerosis by 36% in the aortic arch (p-value = 0.4764, Figure 2B). HL-E97G has a more powerful anti-atherogenic effect, its expression reducing the development of atherosclerosis in the aortic arch by 65% (p-value = 0.0028, Figure 2B). Nevertheless, HL-E97G/evinacumab co-treatment was highly effective, reducing atherosclerotic plaques by 80% (p-value < 0.0001, Figure 2B).

## Discussion

Our team recently identified the second genetic cause associated with familial combined hypocholesterolemia, characterized by very low LDL-C and HDL-C concentrations: the LIPC-E97G gain-of-function variant (Dijk et al. 2022). The E97G mutation is localized in the lid of hepatic lipase (Dijk et al. 2022), where the substrate enters its catalytic pocket. Knowing that this can alter the substrate specificity of the protein (Dugi, Dichek, Santamarina-Fojo 1995), we studied phospholipid lipase and triglyceride lipase activities in several models. While the mutation does not affect triglyceride lipase activity, it does greatly increase the protein’s phospholipase activity (Dijk et al. 2022). This result is the only argument, to date, for the hypocholesterolemic mode of action of HL-E97G. Indeed, the mutant does not appear to internalize cholesterol in the liver nor eliminate it in the feces (Dijk et al. 2022; Sotin et al. 2025). The hypocholesterolemic mode of action of HL-E97G thus remains unclear to this day.

Despite his hypocholesterolemia, the propositus had cardiovascular disease, including developed atherosclerosis, and type 2 diabetes (Dijk et al. 2022). We also know that the atherogenic properties of hepatic lipase are controversial in the literature. However, we have studied the pro- or anti-atherogenic qualities of HL-E97G in several models. Our data show that gain-of-function expression reduces atherosclerotic lesion area in the aortic arch by 97% in humanized APOE3.CETP.Leiden mice and by 41% in dyslipidemic LDLR KO mice (Sotin et al. 2025). These data confirm the anti-atherogenic properties of HL-E97G but, above all, demonstrate a mode of action independent of the LDLR. Nevertheless, the majority of cholesterol-lowering treatments on the market today are LDLR-dependent. They are therefore not very effective for patients with severe hypercholesterolemia, generally due to a defective LDLR, and offer little protection against cardiovascular disease. HL-E97G thus has great therapeutic potential, particularly for familial hypercholesterolemia.

A cholesterol-lowering treatment with an LDLR-independent mode of action has recently been launched: evinacumab, an anti-ANGPTL3 antibody. Loss-of-function variants in the ANGPTL3 gene have been associated, for the first time, with familial combined hypocholesterolemia (Musunuru et al. 2010). This protein stops the hydrolysis activity of endothelial lipase (Shimamura et al. 2007) and lipoprotein lipase (Lee et al. 2009). When ANGPTL3 is blocked, the activity of these two lipases is released. They can then hydrolyze circulating LDL and HDL, inducing hypocholesterolemia. Evinacumab is therefore the most effective treatment, with a 56% reduction in LDL-C in HoFH patients, but none of these patients reach their LDL goal (Béliard et al. 2024). Moreover, evinacumab is ineffective on LDL-C levels when the patient has hypertriglyceridemia (Rosenson et al. 2023). However, we have tested the efficacy of evinacumab/HL-E97G co-treatment in dyslipidemic LDLR KO mice. We still demonstrate very high cholesterol clearance with our mutated protein (−79%), which makes it difficult to see the hypocholesterolemic effect of co-treatment. On the other hand, there is a very nice additive effect. Evinacumab alone reduced lesion area by 36%, HL-E97G by 65%, and co-treatment limited the development of atherosclerosis in the aortic arch by 80%. Our hepatic gain-of-function lipase has strong therapeutic potential for familial hypercholesterolemia but could also be used in combination with evinacumab for hypertriglyceridemia.

## Notes

### Competing Interest Statement

The authors have declared no competing interest.

